# Mitochondrial Oxygen Consumption Drives Lung Tumor Hypoxia and Resistance to Therapy via Copy Number Alteration in Mitochondrial Electron Transport Subunit NDUFB5

**DOI:** 10.64898/2026.07.27.741033

**Authors:** Martin Benej, Katarina Benejova, Adel Fergatova, Rebecca Lisi, Kristina Travis, McKenzie Kreamer, Amy Webb, Caroline Dravillas, Rebecca Hoyd, Esin Bayrali- Ulker, Aashrith Sai Thoutham, Daniel Spakowicz, Nicholas C. Denko

## Abstract

Decades of research have shown that tumor hypoxia is associated with resistance to anti-cancer treatments. Analysis of TCGA gene expression profiles indicates that NSCLC is among the most hypoxic of cancers despite the high levels of oxygen in the surrounding lung tissue. Several groups have shown that extrinsic factors such as poorly formed tumor vascular contributes to tumor hypoxia. Here, we have investigated the possibility that genetic abnormalities within the tumor also contribute to the development of hypoxia. Our analysis of NSCLC patient datasets in the Cancer Genome Atlas (TCGA) PanCancer and ORIEN datasets revealed a strong correlation between tumor hypoxia and amplification of chromosome 3q which is found in up to 40% of NSCLC. Several oncogenic driver genes have been identified in 3q, and we identified a passenger gene encoding mitochondrial complex I subunit NDUFB5 at 3q26.33. To provide experimental evidence that NDUFB5 amplification can drive tumor hypoxia, we have used CRISPR activation technology to generate murine cells overexpressing the endogenous NDUFB5 gene. We found that cells overexpressing NDUFB5 have elevated rates of oxygen consumption, and tumors grown from these cells have increased amounts of hypoxia with associated treatment resistance. Here, we investigate the impact of manipulating NDUFB5 gene expression on mitochondrial complex I activity and experimentally validate the clinical observations that NDUFB5 overexpression leads to increased levels of intratumoral hypoxia and increased resistance to radiation therapy and immunotherapy.

## INTRODUCTION

Tumor hypoxia is a common component of the tumor microenvironment (TME) contributing to resistance to radiation therapy (RT) and immunotherapy in many solid cancers (1–4). Tumor hypoxia exists as an oxygen supply and demand mismatch where the oxygen delivered from the blood cannot meet the metabolic demand of the tumor cells. Oxygen supply is often limited by the well characterized, disorganized tumor vasculature. However, the relationship between the rate of oxygen consumption by the tumor cells and hypoxia has not been reported as a potential contributor of tumor hypoxia. In this work, we ask if we can identify genetic drivers of oxygen consumption that increases tumor hypoxia and therapeutic resistance using publicly available gene expression databases.

Mitochondria are recognized as the major sink for oxygen within the cell. These organelles couple energy production with the consumption of molecular oxygen at cytochrome oxidase at the termination of the electron transport chain (5). As such, mitochondria account for up to 90% of cellular demand for O_2_. The rest of the oxygen demand comes from the use of molecular oxygen as a substrate for enzymes such as oxidases, demethylases, desaturases, and as a terminal electron acceptor for reactions such as disulphide bond formation. Maintaining high mitochondrial function may on one hand provide an advantage for the rapidly proliferating cancer cells by providing both energy and intermediate metabolites. These metabolites are used for de novo synthesis of lipids, amino acids, nucleotides, and buffering of redox stress (6). They are also the source of intermediates such as acetyl-CoA and 2-oxoglutarate that is essential for histone and protein modifications. In hypoxia, mitochondria adapt to reduced oxygen as a major substrate, but continue to produce aspartate and nucleotides and the reducing equivalents that are necessary for pyrimidine synthesis (7, 8). These functions contribute to cancer cell robustness and de novo synthesis allows for cell growth when metabolites are not available in the environment (9). However, the benefit may come at a price – maintaining enhanced mitochondrial function to support proliferation while oxygen is scarce may further decrease the already depleted intratumoral oxygen levels.

To identify an interrelationship between mitochondria oxygen demand and hypoxia, we have looked in the TCGA database for correlative gene expression between components of the mitochondrial electron transport chain and evidence of hypoxia using the Buffa hypoxic gene expressi9on signature. Mammalian complex I is a massive complex consisting of 14 core subunits (those necessary for prokaryotic energy transduction) and 31 supernumerary subunits, all but seven are encoded in nuclear genes (10, 11). We found that NDFUB5, a single subunit of mitochondrial complex I, is frequently amplified in NSCLC and in tumors where it is amplified, there is increased hypoxia (increased Buffa score). NDUFB5 is one of the supernumerary subunits embedded in the inner mitochondrial membrane, spanning almost the entire membrane arm of complex 1. It is thought to play an essential role in complex I assembly by co-joining the proximal and distal modules together (10). Interestingly, NDUFB5 does not have any recognized enzymatic function and is not directly involved in passing electrons or pumping protons (11). We experimentally demonstrate that overexpression of NDFUB5 elevates complex I activity and mitochondrial oxygen consumption leading to increased hypoxia levels and treatment resistance in mouse models of NSCLC.

## RESULTS

### Mitochondrial subunit gene expression associates with high levels of hypoxia in NSCLC

Significant scientific attention has been focused on studying how oncogenic mutations in genes such as epidermal growth factor receptor (EGFR), K-ras (KRAS) and anaplastic lymphoma kinase (ALK) promote progression and treatment resistance in NSCLC (12–15). We decided to take an alternative approach and work backward from the therapy resistant hypoxic phenotype to identify genetic drivers of hypoxia. We first used the Buffa hypoxic gene expression profile to categorize tumors in the TCGA PanCan database. We calculated the Buffa score for each tumor as a way to quantify hypoxia and plotted the values for each type of cancer. **Fig. 1a** shows the raincloud plot of Buffa scores and how these values vary from cancer types displaying the least hypoxia (top, thyroid) to the most hypoxia (bottom, squamous cancers of head and neck, cervix and lung). To put these values in context, we next calculated the Buffa scores for the corresponding non-cancerous tissues from the GTEx database (16), and the samples from the normal adjacent tissue in the TCGA PanCan (17). Comparison of each normal tissue and cancer is in **Fig. S1A,** and shows that for each tissue type the tumors have higher buffa scores, indicating quantitatively more hypoxia. Samples for NSCLC are shown in **Fig. 1B** and clearly show that even when the normal adjacent tissue is well oxygenated, the lung cancers are very hypoxic, with squamous cell tumors showing more hypoxia than tumors with adenocarcinoma histology.

**Fig. 1.**
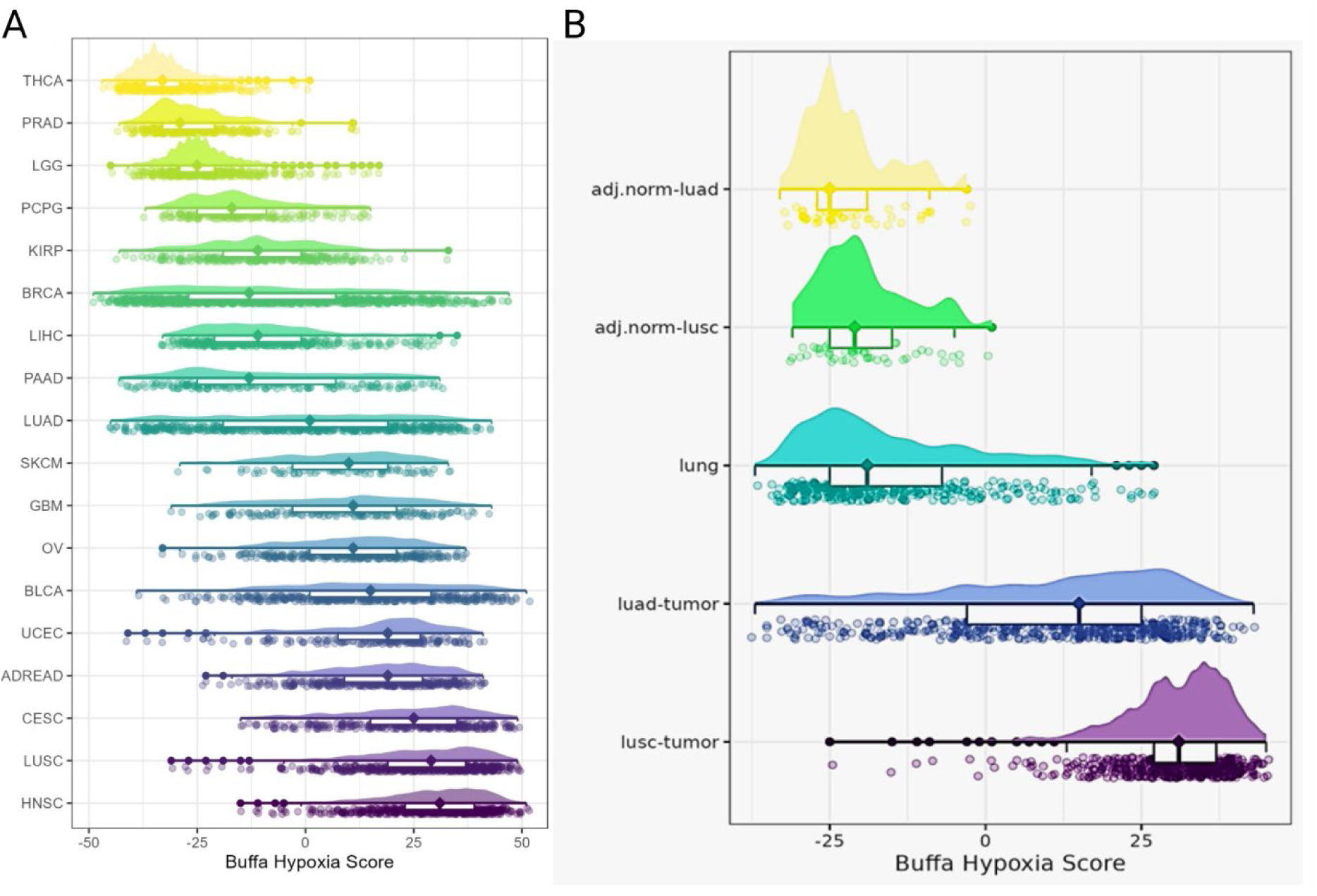
Tumor hypoxia in TCGA samples. **(A)** Raincloud plot of Buffa hypoxia gene expression scores for 18 tumor types of the PanCan dataset ordered from least hypoxic in yellow at the top to most hypoxic in purple at the bottom. **(B)** Normal lung (GTEx), normal adjacent, lung adeno and lung squamous tumors (TCGA).

We next looked for relationships between Buffa scores and genes that may be driving the hypoxia. We found that in lung cancers, activating mutations in the genetic drivers KRAS, EGFR and ALK were just as common in Buffa low as Buffa high tumors, indicating that these mutations were not responsible for hypoxia and its associated therapy resistance. (**Fig. S1B, C**). Other investigators have clearly shown that hypoxia is a powerful stimulus for angiogenesis and many angiogenic growth factors, and their receptors are hypoxia inducible such as VEGFs/VEGFRs. Instead of investigating the role of hypoxia on oxygen supply through angiogenesis, we reasoned that the mitochondria are the major sink for oxygen in the cell, and decided to investigate the possibility that high levels of mitochondrial gene expression could lead to high oxygen demand within the tumor and elevated hypoxia. We therefore generated an “oxygen consumption rate score” (OCR score) that combined the expression of all 91 nuclear genes encoding proteins in the mitochondrial electron transport chain. We calculated this score for each tumor and then compared the OCR score with the Buffa score for all the tumors in TCGA Pancan database. Interestingly, we found a significant positive relationship in 16 of 18 tumor types (**Fig. S2A**). This relationship indicates that high expression of mitochondrial ETC genes and subsequent oxygen demand could contribute to hypoxia in tumors.

Therefore, we individually examined the level of expression of each of the 91 genes of the OCR score (**Fig. 2A**) with respect to the Buffa score. Several genes’ expressions were positively related to Buffa score such as NDUFV1, NDUFA9 and SDHA, but by far the strongest correlation was with NDUFB5 (**Fig. 2B**). To better understand the variable expression of NDUFB5, we examined the copy number variation and found that NDUFB5 mRNA level was highly related to gene copy number, with over half the tumors showing some level of NDUFB5 amplification (**Fig. 2C**). NDUFB5 is located on chromosome 3q:26.33 and is frequently amplified in NSCLC with other regions of 3q, which is reported to occur in up to 40% of lung cancers (18). We hypothesized that NDUFB5 is not necessarily an oncogenic driver, but rather a passenger gene in a region close to any oncogenic drivers (such as SOX2, PIK3CA, or PRKCI) that are selected for when 3q is amplified. Finally, we analyzed the relationship between NDUFB5 mRNA expression and Buffa score for each of the NSCLC in TCGA. There is a highly significant and sensitive positive relationship between the two values (**Fig. 2D**).

**Fig. 2.**
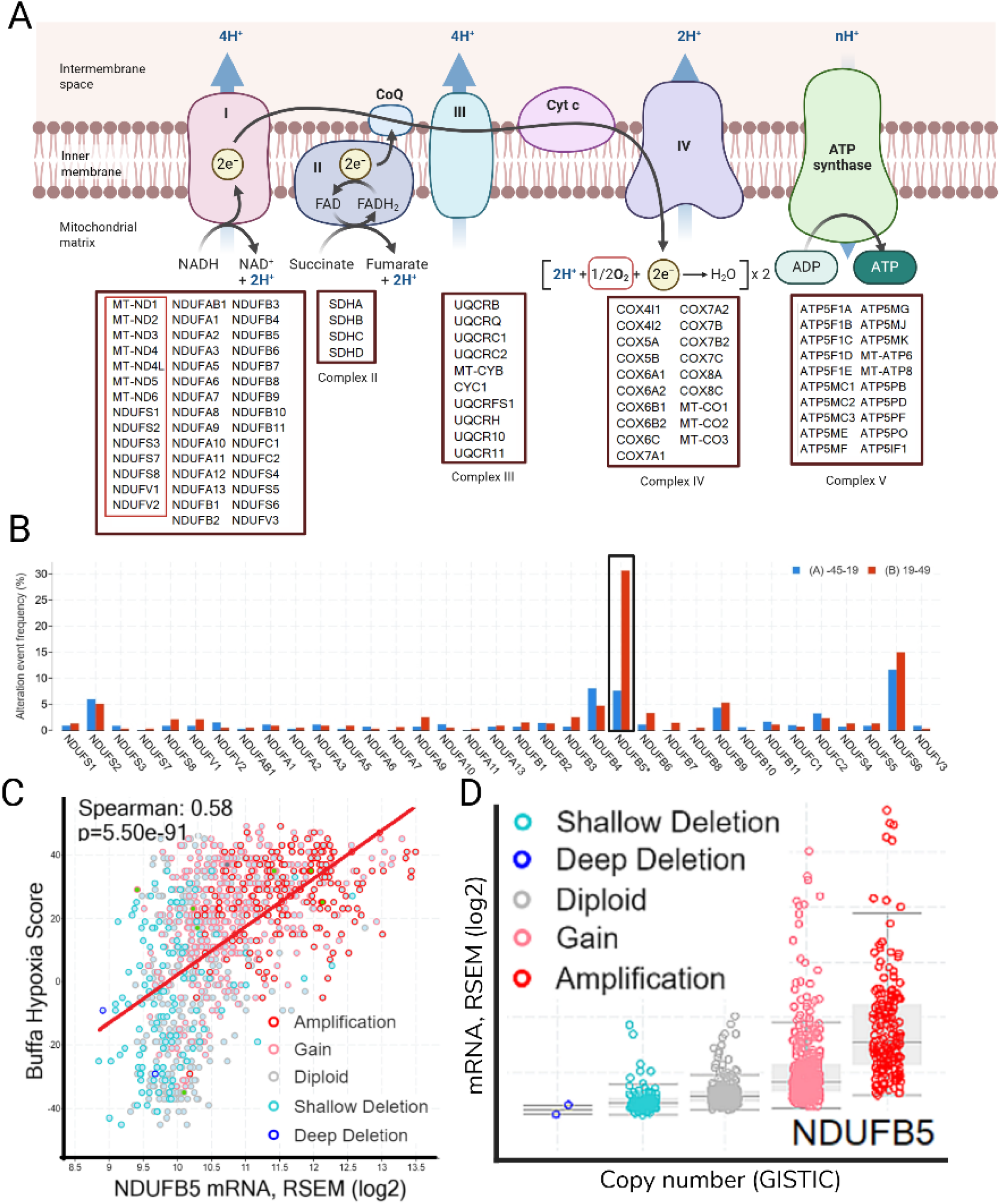
NDUFB5 mRNA expression relates to Buffa hypoxia score. **(A)** The role of molecular oxygen as the final electron acceptor in the process of mitochondrial respiration. 91 genes of the OCR score shown that encode mitochondrial complex I-V subunits are shown. **(B)** TCGA analysis of gene expression of complex 1 genes in lung cancer separated by oxic (*blue bars*) vs hypoxic (*red bars*) tumors. Median split of Buffa hypoxia score was used to stratify patients into oxic vs hypoxic tumors. **(C)** NDUFB5 mRNA expression in patients with amplification (> 5 copies; red); moderate gain of copies (3-5 copies; *pink*); diploid (2 copies, *gray*); and deletion (< 2 copies, *light blue*) (n=1,053). **(D)** TCGA analysis of NDUFB5 copy number against respective mRNA levels in NSCLC tumor samples.

### Experimental evidence for elevated mitochondrial activity after NDUFB5 amplification

We next experimentally confirmed the mechanistic connection between NDUFB5 overexpression leading to elevated mitochondrial oxygen consumption and model tumor hypoxia. We engineered the murine lung adenocarcinoma CMT167 and squamous cell carcinoma cell KLN205 to overexpress NDUFB5 from the endogenous locus using CRISPR activation constructs (Santa Cruz Biotechnology). Individual overexpressing clones were confirmed by western blot and pooled together for Seahorse XF96 analysis of antimycin sensitive mitochondrial OCR (**Fig. 3A, S3A**). We found that a 2–3-fold increase in NDUFB5 expression results in a 50-80% increase in baseline mitochondrial oxygen consumption rate by Seahorse XF analysis (**Fig. 3B**). To test the hypothesis that elevated mitochondrial OCR drives intratumoral hypoxia, we grew CMT167 or KLN 205 Parent or NDUFB5 OE cells as tumors in the flanks of immunocompetent mice (C57Bl6 or DBA/2 host mice respectively). When tumors reached 500-800 mm^3^, we injected the mice with pimonidazole an hour before harvesting the tumors to mark the hypoxic regions. Pimonidazole is a 2-nitroimidazole that is reduced intracellularly and makes covalent adducts in hypoxic regions that can be detected by monoclonal antibody. Sections of tumors were cut and stained for pimonidazole adducts and intratumoral hypoxia levels were quantified using immunofluorescence. **Fig. 3C** **and S3B** show that pimonidazole staining was significantly elevated in all OE tumors compared to parent tumors, supporting the model that NDUFB5 expression can drive intratumoral hypoxia. To evaluate the mechanism by which NDUFB5 overexpression elevates mitochondrial OCR, we compared the rate of ETC complex I activity by measuring the rate of NADH-dependent oxidation of a synthetic ubiquinone in isolated mitochondria from parent and OE cells and observed a significant increase in biochemical rate of complex 1 activity in the overexpressing CMT167 cells (**Fig. 3D**). Interestingly, we found that these cells have a moderate increase of mitochondrial mass as determined by nonyl acridine orange (NAO) staining and FACS, and no significant change in membrane potential or mitochondrial ROS production (**Fig. 3E**). Finally, previous studies established that mitochondrial morphology can play an important role in regulating the mitochondrial function (19). To evaluate whether NDUFB5 overexpression affects mitochondrial morphology, we compared the mitochondria of CMT167 parent or NDUFB5 overexpressing cells by transmission electron microscopy. We determined that compared to the parent cells, mitochondria in NDUFB5 overexpressing cells have a significantly narrower shape with more cristae per mitochondrion (**Fig. 3G-H**), which other groups have associate with increased mitochondrial function (19).

**Fig. 3.**
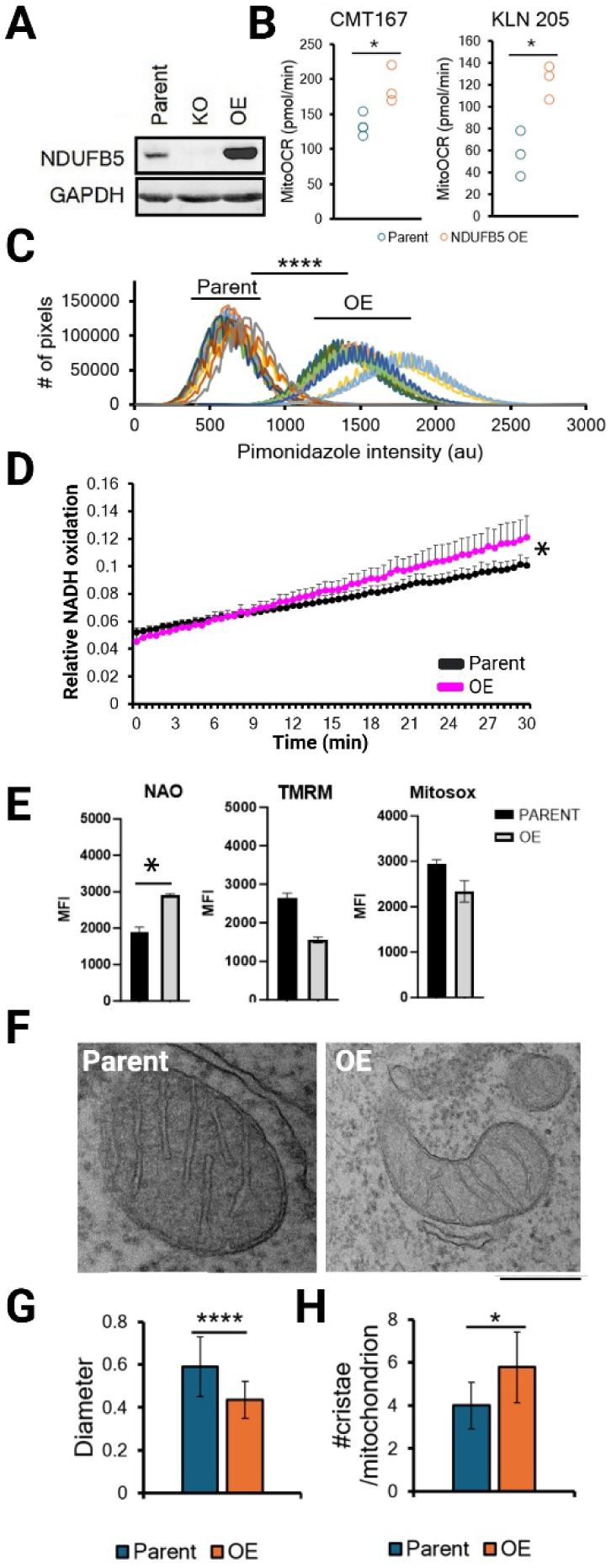
**Experimental confirmation of elevated OCR and tumor hypoxia after NDUFB5 over expression**. (**A**) Representative western blot validation of CRISPR/cas9 mediated knockout (KO) or CRISPRa-mediated overexpression (OE) of endogenous NDUFB5 in CMT167 cell line. (**B**) Seahorse analysis showing elevated levels of mitochondrial oxygen consumption (mitoOCR) in CMT167 or KLN 205 NDUFB5 overexpressing cells. (**C**) Quantification of hypoxic fractions in CMT167 parent and OE tumors grown in C57BL/6 mice (n=4). Values are histograms of arbitrary units of pimonidazole intensity in IF slides against number of pixels for each intensity value, quantified from 10 images per animal. P value was calculated with t test. **(D)** The ability of mitochondrial complex I to oxidize NADH was measured using spectrophotometric assay as described in (20). **(E)** Evaluation of mitochondrial mass (NAO, nonyl acridine orange); membrane potential (TMRM, tetramethylrhodamine); or ROS production (MitoSOX) in parent or NDUFB5 overexpressing CMT167 cells by flow cytometry. **(F)** Representative transmission electron microscopy (TEM) images of mitochodnria in CMT167 Parent and OE cells. Line represents 200 nm. **(G-H)** Quantification of mitochondrial diameter (G) and cristae/mitochondrion (H) in CMT167 Parent vs OE cells; n=30 mitochondria per group. P value was calculated with t test. * < 0.05; **** < 0.0001.

### NDUFB5 expression correlates with therapeutic resistance in NSCLC patients

Hypoxia is a recognized barrier to effective radiation therapy and immunotherapy (1, 2, 4, 21). We first validated that Buffa hypoxia score predicts patient survival in a cohort of OSU patients with NSCLC treated with stereotactic body radiation therapy (SBRT, n=219) or first-line immune checkpoint blockade (ICB) (n=85), available through the OSU Total Cancer Care® Protocol (**Fig. 4A-B**). This is consistent with our previously published series (22). Because we established that increased NDUFB5 mRNA expression correlates with increased Buffa scores, we next evaluated whether NDUFB5 mRNA expression itself can predict patient outcomes within the TCC datasets. Interestingly, in NSCLC patients discriminated by median NDUFB5 mRNA expression, we observed a significant decrease of overall survival in the NDUFB5-high cohort (n=84, p=0.0187) (**Fig. 4C**). Moreover, if we stratify the patients who received ICB by NDUFB5 mRNA, there was a trend to reduced survival for the over expressing cohort. However, if we analyze the SU2C MARK lung cancer database (23), we do see a significantly worse prognosis for patients with amplified NDUFB5 if they are treated with primary ICB (**Fig. 4D).** This dataset also has the tumor mutational burden (TMB) for these patients. If we use TMB to further stratify patients NDUFB5 amplified versus diploid, we find that high TMB predicted better outcome for the NDUFB5 diploid patients, but did not impact outcomes in the NDUFB5 amplified patients, suggesting hypoxia is dominant over TMB as a regulator of patient response to ICB (**Fig. S4A**). To investigate possible mechanisms for this effect, we used CIBERSORTx-based analysis of deconvoluted TCGA patient data, showing a 50% decrease of absolute CD8^+^ T cell abundance in patients with more than the diploid number of NDUFB5 copies (**Fig. S4B**). These findings support the model that elevated NDUFB5 expression is associated with increased tumor hypoxia, leading to decreased patient survival after either radiotherapy or immunotherapy.

**Fig. 4.**
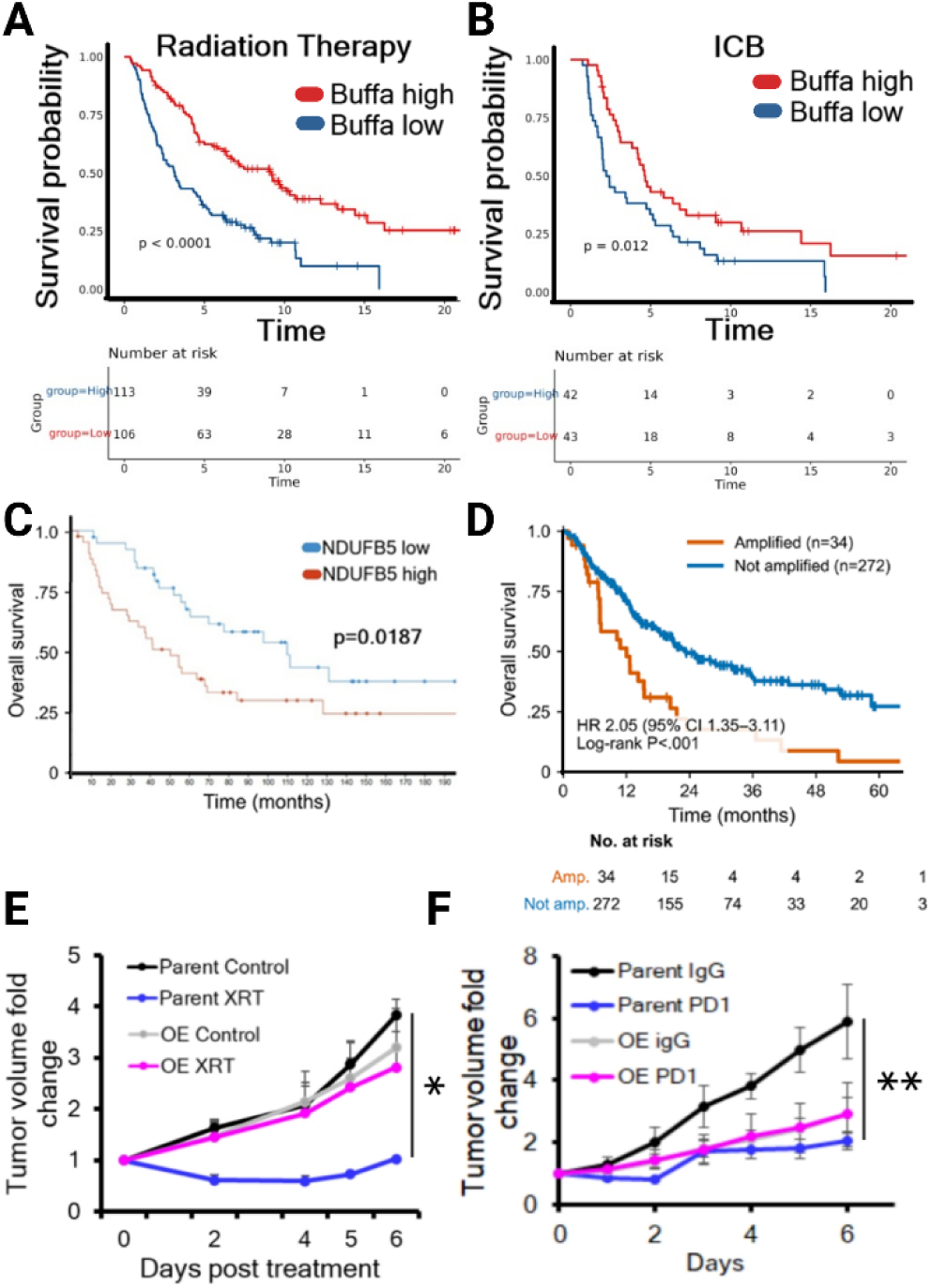
(A-B) Overall survival in NSCLC patients treated at OSU with radiation therapy (A, n=219) or first-line immune checkpoint blockade (B, n=85). Median split of Buffa hypoxia score was used to stratify patients into oxic vs hypoxic tumors. **(C)** Overall survival in OSU NSCLC patients treated with radiation therapy. Median split of NDUFB5 mRNA was used to stratify patients into NDUFB5 high (*red*) vs NDUFB5 low (*blue*); (n=84). **(D)** Correlation between poor overall survival and NDUFB5 mRNA expression in OSU NSCLC patients treated with immune checkpoint blockade (n=85). **(E-F)** Quantification of tumor growth delay of CMT167 parent and NDUFB5 overexpressing tumors grown in C57BL/6 mice. The mice received either a single 8 Gy dose using the Small Animal Radiation Research Platform (E); or three cycles of either rat isotype IgG control or anti-PD-1 ICI antibody (100 μg I.P.) on Day 0, 3 and 5. Curves represent mean tumor volumes ± SEM. P values were calculated for PD-1 vs IgG in each group (parent or OE) by t test. *P < 0.05; **P < 0.01.

To validate these clinical findings experimentally, we next compared the tumors from CMT167 or KLN205 cells overexpressing NDUFB5 to parental tumors for response to either radiotherapy or immune checkpoint blockade. Radiation therapy was delivered via a single 8 Gy dose using image guidance on the Small Animal Radiation Research Platform (SARRP); immunotherapy was delivered via a single dose of anti-PD-1 antibody (100 μg, i.p. against rat IgG isotype control, BioXcell). We observed that for both treatment modalities, the NDUFB5 overexpressing tumors were highly resistant when compared to the parent tumors (**Fig. 4E-4F, S3C**). We confirmed that the cells did not have cell autonomous resistance to radiotherapy because WT and NDUFB5 overexpressing cells irradiated in vitro showed identical cell kill. One mechanism by which hypoxia leads to immune privilege is through exclusion of effector CD8 T cells (2). To evaluate whether NDUFB5 overexpression causes this in our model tumors, we sectioned naïve tumors and quantified the infiltrating immune cells by immunofluorescence using anti-CD8 antibody. We found a significant decrease of CD8^+^ T cell abundance in the NDUFB5 overexpressing tumors (**Fig.S4 C-D**).

## DISCUSSION

In this manuscript, we report that increased copy number and mRNA expression of a single, supernumerary, structural mitochondrial complex I subunit is sufficient to increase mitochondrial OCR in cells, increase hypoxia in model tumors, and causes tumor resistance to radio- and immuno-therapies in model NSCLC.

NSCLC is categorized as primarily either of adenocarcinoma or squamous cell carcinoma histology. Squamous variants are typically more hypoxic and this may in part explain the higher rate of local recurrence post SBRT (22, 24), and inferior ICB outcomes compared to adenocarcinomas (25, 26). Incidentally, squamous cell carcinomas are also associated with elevated NDUFB5 mRNA expression and copy number; some publicly available datasets show that up to 70% of NSCLC with squamous histology have more than two copies of NDUFB5. This subtype has been shown to harbor an amplification of 3q, centered around the 30Mb 3q26-3q28 containing the NDUFB5 locus (18). Fine mapping of this amplicon has identified several candidate genes that could be contributing to oncogenic transformation at this locus including PIK3CA, TP73L, EIF4G and PKC iota. Future work will investigate whether these loci are passengers or whether their co-amplification contributes to the high rates of hypoxia observed in NDUFB5-high tumors.

The hypoxic tumor microenvironment is a known driver of metastatic spread(27) at least in part by actively promoting selection for the most aggressive cancer cells able to thrive in highly acidic conditions and oxygen scarcity. One of the possible implications of NDUFB5 overexpression-driven mitochondrial oxygen demand is therefore an increased rate of metastatic spread. Moreover, previous studies in kidney cancer reported that increased mitochondrial complex I activity is associated with increased metastasis formation (28). Future mechanistic directions will explore whether biopsy specimens from distant metastases show increased levels of NDUFB5 expression which will elucidate one of the mechanisms of disease progression in NDUFB5-high NSCLCs.

To date, NDUFB5 has been considered a supernumerary complex I subunit without enzymatic activity (10). It is intriguing that overexpression of a single subunit, that is not one of the fourteen core complex I subunits, leads to increased complex I activity. We have not observed a change in mRNA or protein levels of other complex I subunits upon overexpressing NDUFB5, nor have we observed a change in mitochondrial mass or membrane potential. In the fully assembled complex I, NDUFB5 spans the transmembrane arm of the complex and is required for correct subassembly of the transmembrane domain modules of complex I (11). Mitochondrial complex I is one of three complexes (complexes I, III and IV) where the electron flow is coupled to the pumping of protons across the inner mitochondrial membrane, creating a proton gradient that drives ATP synthase (29). Because the elements of the proton-translocating machinery are contained mostly within the transmembrane domain of the complex, modified expression of NDUFB5 may alter the proton pumping capacity thereby affecting the proton flux into the intermembrane space resulting in changes of ATP synthesis and membrane potential. Furthermore, mitochondrial complex I was shown to assemble into a multimeric supercomplex with complex III (ubiquinol-cytochrome *c* oxidoreductase) and complex IV (cytochrome *c* oxidase) (30, 31), which can be enhanced in hypoxia (32). The supercomplex formation leads to reduced electron leak and increases ETC efficiency (33). Because NDUFB5 facilitates proper assembly of the transmembrane domain modules and complex I binds to complex III via the distal region of the transmembrane domain (34), altered NDUFB5 expression levels may affect the assembly state of complex I (free or bound to complex III as a part of the supercomplex). The exact mechanistic nature of the NDUFB5-driven increase of mitochondrial oxygen demand therefore remains to be elucidated in future studies and will contribute to better understanding of complex I biology and its impact on NSCLC progression and provides the potential for an additional marker for predicting treatment response in patients.

## ACKNOWLEDGEMENTS

This work was supported by funding from NCI grants R03 CA280489 (MB) and K22 CA282363 (MB); and NCI grants R01 CA255344 (ND) and R01 CA262388 (ND). The authors would like to acknowledge assistance from the OSUCCC Small Animal Imaging Shared Resource and Campus Microscopy and Imaging Facility Shared Resource; and from the Total Cancer Care (TCC) program, which is part of the Center’s Biospecimen Services Shared Resource (BSSR).

## METHODS

### Cell Lines

All cell lines were purchased from the American Type Culture Collection (ATCC) and grown in DMEM (Corning) supplemented with 10% FBS (Seradigm) and 1% Pen/Strep (Fisher Bioreagents). Cell counts and cellular proliferation were established using automated cell counter to determine the cell number and trypan blue exclusion assay to establish the fraction of viable/dead cells. All cell lines were authenticated using short tandem repeat (STR) profiling within the past 3 years. All experiments were performed with mycoplasma-free cells. NDUFB5 CRISPR activation plasmids were obtained from Sant Cruz Biotechnology (sc-425783-ACT) and CMT167 or KLN205 cell lines were transfected using Lipofectamine 2000 according to the manufacturer’s protocol (Thermo Fisher Scientific). After selection in 4 μg/mL Blasticidin S HCL, 300 μg/mL Hygromycin B and Puromycin 5 μg/mL (Santa Cruz) the cells were diluted into single-cell suspensions, and individual clones were screened by western blot.

### Western Blotting

Proteins were extracted with 1% v/v Triton X-100, 0.5% w/v NP-40, 150 mM NaCl, and 50 mM Tris (pH 7.5), quantified using the BCA Kit (Thermo Scientific, Waltham, MA, USA), and separated on 10% SDS-PAGE under reducing conditions. Proteins of interest were detected using antibodies against firefly NDUFB5 and β-actin (Santa Cruz Biotech, Dallas, TX, USA). All replicates of the individual experiments were analyzed from the respective run with appropriate loading controls.

### Seahorse Analysis of the OCR

Oxygen consumption rate (OCR) was measured using Seahorse XF96 (Agilent Technologies). The cells were seeded overnight, washed with prewarmed XF Calibrant, and replaced with unbuffered Assay Medium (pH 7.4, 5 mM glucose, 1 mM l-glutamine) and incubated 2 h.

### Animal experiments

The CMT167 cells parent or NDUFB5 overexpressing cells (1 × 10^6^) were injected s.c. into the flanks of 7-wk-old female C57BL/6; or the KLN205 parent or NDFUB5 overexpressing cells (2 × 10^6^) were injected into the flanks of 7-wk-old female Dba/2 mice following Institutional Animal Care and Use Committee (IACUC)-approved protocols. Caliper measurements of opposing diameter were used to calculate the tumor volumes. Pimonidazole adducts were visualized in hypoxic regions within histological sections of tumor tissues (39). Mice bearing CMT167 or KLN205 parent or NDUFB5 overexpressing tumors were treated with 60 mg/kg pimonidazole i.p., and tumors were harvested at 90 min. Frozen sections were stained with anti-pimonidazole rabbit antibody and anti-rabbit Alexa Fluor 488. The hypoxic fraction of each tumor was quantified by thresholding signal at 50% of the maximum signal on control sections. The area covered by pimonidazole-positive cells was evaluated from 20 images per animal and averaged. For RT response evaluation, upon reaching 150 mm^3^, the tumors were visualized by cone beam CT, and treatment plans were calculated using SARRP software. X-rays were delivered with a single beam delivering 8 Gy using the Small Animal Research Radiation Platform (SARRP; Xstrahl). For the immunotherapy response evaluation, upon reaching 150 mm^3^, the mice received three doses of anti-PD-1 antibody (100 μg, i.p. against rat IgG isotype control, BioXcell). Tumor volumes were measured until the posttreatment volume increased threefold.

### Cell lysate preparation for Complex I Assay

Cells were treated with 0.1 µg/mL doxycycline for 48 h, after which they were harvested, washed with PBS, and counted. Cells were adjusted to a density of 5 × 10⁶ cells per sample and pelleted by centrifugation at 1000 rpm for 5 min, followed by one freeze-thaw cycle at −80 °C. Pelleted cells were resuspended in 750 µL of ice-cold SET lysis buffer (0.25M sucrose, 2 mM EDTA, 10 mM Tris, pH 7.4) and incubated on ice for 5–10 min before homogenization using a pre-chilled Dounce homogenizer. The number of strokes was optimized for each cell line to achieve efficient cell lysis, with lysis efficiency assessed by trypan blue exclusion. The resulting whole-cell lysate was used directly in the Complex I activity assay.

### Spectrophotometric Complex I Assay

Complex I activity assay was adapted from Janssen et al. (2007). Complex I activity was assessed by monitoring 2,6-dichloroindophenol (DCPIP) reduction at 600 nm, reflecting electron transfer from decylubiquinone to the dye.

The reaction mixture contained 25 mM potassium phosphate, 3.5 g/L BSA, 60 µM DCPIP, 70 µM decylubiquinone, 1.0 µM antimycin-A (pH 7.8), in a final volume of 100 uL. BSA was essential for solubilizing decylubiquinone and rotenone in this assay.

The reaction was performed in a total volume of 100 µL containing 25 mM phosphate buffer (pH 7.2), 3.5 mg/mL BSA, 60 µM DCPIP, 70 µM decylubiquinone, 10 µM antimycin, and 10 µL of cell lysate. Samples were preincubated with either 1 uM rotenone, 10 µM papaverine, or an equivalent volume of vehicle (control) for 5 min at room temperature. 0.2mM NADH was added to initiate the reaction, and absorbance at 600 nm was recorded over 20 min, with the rate of absorbance decline reflecting Complex I activity.

Protein concentration in 10 µL lysate aliquot was determined by bicinchoninic acid (BCA) assay, and Complex I activity values were normalized using the protein concentration of each corresponding sample.

## REFERENCES

1. Baginska J, Viry E, Paggetti J, Medves S, Berchem G, Moussay E, et al. The critical role of the tumor microenvironment in shaping natural killer cell-mediated anti-tumor immunity. Front Immunol. 2013;4:490.

2. Lee CT, Mace T, Repasky EA. Hypoxia-driven immunosuppression: a new reason to use thermal therapy in the treatment of cancer? Int J Hyperthermia. 2010;26(3):232–46.

3. Liu Y-N, Yang J-F, Huang D-J, Ni H-H, Zhang C-X, Zhang L, et al. Hypoxia Induces Mitochondrial Defect That Promotes T Cell Exhaustion in Tumor Microenvironment Through MYC-Regulated Pathways. Frontiers in Immunology. 2020;11.

4. Thomlinson RH, Gray LH. The histological structure of some human lung cancers and the possible implications for radiotherapy. Br J Cancer. 1955;9(4):539–49.

5. Niethammer P, Kueh HY, Mitchison TJ. Spatial patterning of metabolism by mitochondria, oxygen, and energy sinks in a model cytoplasm. Curr Biol. 2008;18(8):586–91.

6. Benej M, Papandreou I, Denko NC. Hypoxic adaptation of mitochondria and its impact on tumor cell function. Semin Cancer Biol. 2024;100:28–38.

7. Martinez-Reyes I, Chandel NS. Mitochondrial TCA cycle metabolites control physiology and disease. Nat Commun. 2020;11(1):102.

8. Garcia-Bermudez J, Baudrier L, La K, Zhu XG, Fidelin J, Sviderskiy VO, et al. Aspartate is a limiting metabolite for cancer cell proliferation under hypoxia and in tumours. Nat Cell Biol. 2018;20(7):775–81.

9. Weinberg SE, Chandel NS. Targeting mitochondria metabolism for cancer therapy. Nat Chem Biol. 2015;11(1):9–15.

10. Stroud DA, Surgenor EE, Formosa LE, Reljic B, Frazier AE, Dibley MG, et al. Accessory subunits are integral for assembly and function of human mitochondrial complex I. Nature. 2016;538(7623):123–6.

11. Padavannil A, Ayala-Hernandez MG, Castellanos-Silva EA, Letts JA. The Mysterious Multitude: Structural Perspective on the Accessory Subunits of Respiratory Complex I. Front Mol Biosci. 2021;8:798353.

12. da Cunha Santos G, Shepherd FA, Tsao MS. EGFR mutations and lung cancer. Annu Rev Pathol. 2011;6:49–69.

13. Morris SW, Kirstein MN, Valentine MB, Dittmer KG, Shapiro DN, Saltman DL, et al. Fusion of a kinase gene, ALK, to a nucleolar protein gene, NPM, in non-Hodgkin’s lymphoma. Science. 1994;263(5151):1281–4.

14. Sasaki T, Rodig SJ, Chirieac LR, Jänne PA. The biology and treatment of EML4- ALK non-small cell lung cancer. Eur J Cancer. 2010;46(10):1773–80.

15. Cascetta P, Marinello A, Lazzari C, Gregorc V, Planchard D, Bianco R, et al. KRAS in NSCLC: State of the Art and Future Perspectives. Cancers (Basel). 2022;14(21).

16. The GTEx Consortium atlas of genetic regulatory effects across human tissues. Science. 2020; 369(6509):1318–30.

17. Liu J, Lichtenberg T, Hoadley KA, Poisson LM, Lazar AJ, Cherniack AD, et al. An Integrated TCGA Pan-Cancer Clinical Data Resource to Drive High-Quality Survival Outcome Analytics. Cell. 2018;173(2):400–16.e11.

18. Qian J, Massion PP. Role of chromosome 3q amplification in lung cancer. J Thorac Oncol. 2008;3(3):212–5.

19. Galloway CA, Lee H, Yoon Y. Mitochondrial morphology-emerging role in bioenergetics. Free Radic Biol Med. 2012;53(12):2218–28.

20. Janssen AJ, Trijbels FJ, Sengers RC, Smeitink JA, van den Heuvel LP, Wintjes LT, et al. Spectrophotometric Assay for Complex I of the Respiratory Chain in Tissue Samples and Cultured Fibroblasts. Clinical Chemistry. 2007;53(4):729–34.

21. Saunders M, Dische S. Clinical results of hypoxic cell radiosensitisation from hyperbaric oxygen to accelerated radiotherapy, carbogen and nicotinamide. Br J Cancer Suppl. 1996;27:S271–8.

22. Eustace NJ, Sebastian NT, Webb A, Shilo K, Robb R, Amini A, et al. Hypoxic Gene Expression Signature as a Predictor of Recurrence and Mortality in Early-Stage Non-Small Cell Lung Cancer Treated With Stereotactic Body Radiation Therapy. JCO Precis Oncol. 2025;9:e2400659.

23. Ravi A, Hellmann MD, Arniella MB, Holton M, Freeman SS, Naranbhai V, et al. Genomic and transcriptomic analysis of checkpoint blockade response in advanced non-small cell lung cancer. Nat Genet. 2023;55(5):807–19.

24. Kita N, Tomita N, Takaoka T, Sudo S, Tsuzuki Y, Okazaki D, et al. Comparison of Recurrence Patterns between Adenocarcinoma and Squamous Cell Carcinoma after Stereotactic Body Radiotherapy for Early-Stage Lung Cancer. Cancers (Basel). 2023;15(3).

25. Herbst RS, Giaccone G, de Marinis F, Reinmuth N, Vergnenegre A, Barrios CH, et al. Atezolizumab for First-Line Treatment of PD-L1-Selected Patients with NSCLC. N Engl J Med. 2020;383(14):1328–39.

26. Johnson ML, Cho BC, Luft A, Alatorre-Alexander J, Geater SL, Laktionov K, et al. Durvalumab With or Without Tremelimumab in Combination With Chemotherapy as First-Line Therapy for Metastatic Non-Small-Cell Lung Cancer: The Phase III POSEIDON Study. J Clin Oncol. 2023;41(6):1213–27.

27. Le QT, Denko NC, Giaccia AJ. Hypoxic gene expression and metastasis. Cancer Metastasis Rev. 2004;23(3-4):293–310.

28. Bezwada D, Perelli L, Lesner NP, Cai L, Brooks B, Wu Z, et al. Mitochondrial complex I promotes kidney cancer metastasis. Nature. 2024;633(8031):923–31.

29. Galkin A. Water route for proton pumping in mitochondrial complex I. Nat Struct Mol Biol. 2020;27(10):1–3.

30. Schägger H, Pfeiffer K. Supercomplexes in the respiratory chains of yeast and mammalian mitochondria. Embo j. 2000;19(8):1777–83.

31. Guo R, Zong S, Wu M, Gu J, Yang M. Architecture of Human Mitochondrial Respiratory Megacomplex I(2)III(2)IV(2). Cell. 2017;170(6):1247–57.e12.

32. Chen YC, Taylor EB, Dephoure N, Heo JM, Tonhato A, Papandreou I, et al. Identification of a protein mediating respiratory supercomplex stability. Cell Metab. 2012;15(3):348–60.

33. Blaza JN, Serreli R, Jones AJY, Mohammed K, Hirst J. Kinetic evidence against partitioning of the ubiquinone pool and the catalytic relevance of respiratory-chain supercomplexes. Proceedings of the National Academy of Sciences. 2014;111(44):15735–40.

34. Arroum T, Borowski MT, Marx N, Schmelter F, Scholz M, Psathaki OE, et al. Loss of respiratory complex I subunit NDUFB10 affects complex I assembly and supercomplex formation. Biol Chem. 2023;404(5):399–415.

