## Supplementary Data for "Mitochondrial Oxygen Consumption Drives Lung Tumor Hypoxia and Resistance to Therapy via Copy Number Alteration in Mitochondrial Electron Transport Subunit NDUFB5"

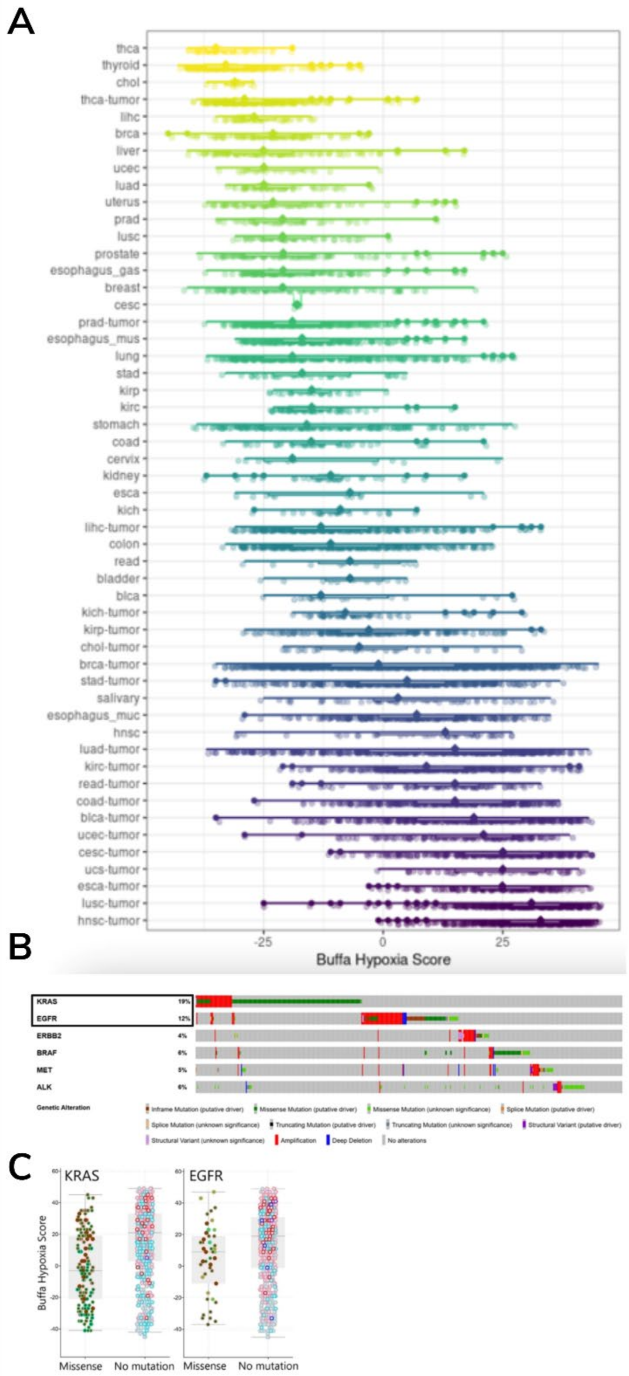


**Fig. S1. (A)** Raincloud plot of Buffa hypoxia gene expression scores in TCGA PanCancer datasets and respective normal tissue from the Gtex database. **(B)** Driver mutation frequency in NSCLC datasets (TCGA). **(C)** KRAS or EGFR mutations do not significantly affect Buffa hypoxia scores.

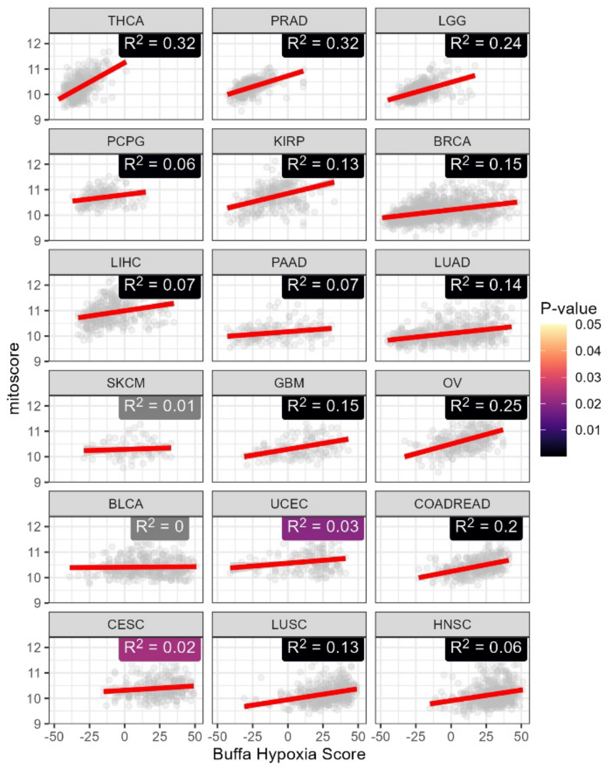


**Fig. S2. (A)** Mitoscore versus Buffa values across the 18 TCGA Pancan datasets showing significant positive statistical correlation in all but skin and bladder cancer. Statistical significance indicated by depth of color for R2.

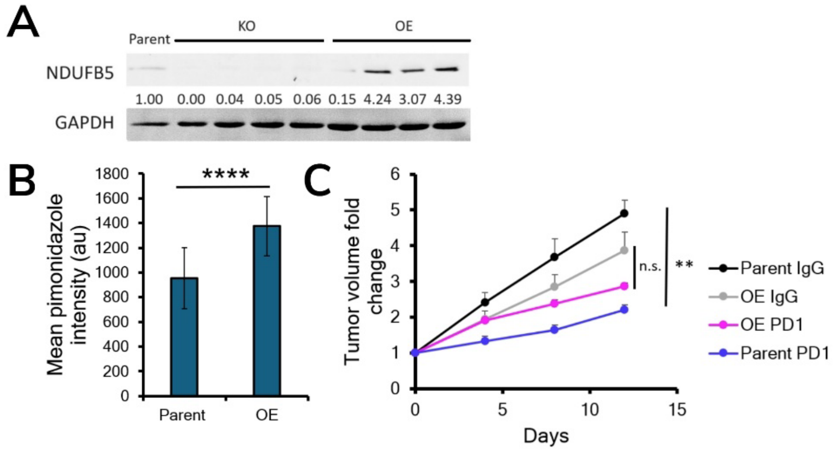


**Fig. S3. (A)** Representative western blot validation of CRISPR/cas9 mediated knockout (KO) or CRISPRa-mediated overexpression (OE) of endogenous NDUFB5 in KLN205 cell line. **(B)** Quantification of hypoxic fractions in KLN205 parent and OE tumors grown in Dba/2 mice (n=4). Values are arbitrary units of pimonidazole intensity in IF slides against number of pixels for each intensity value, quantified from 10 images per animal. P value was calculated with t test. **(C)** Quantification of tumor growth delay of KLN205 parent and NDUFB5 overexpressing tumors grown in Dba/2 mice. The mice received three cycles of either rat isotype IgG control or anti-PD-1 ICI antibody (200 µg I.P.) on Day 0, 4 and 8. Curves represent mean tumor volumes ± SEM. P values were calculated for PD-1 vs IgG in each group (parent or OE) by t test. **P < 0.01.

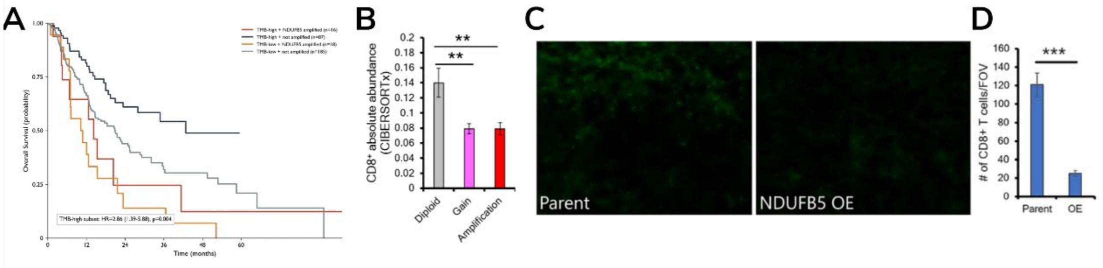


**Fig. S4. (A)** Patients from the SU2C MARK NSCLC database were stratified by NDUF amplification and then further stratified by median tumor mutational burden. Note that TMB only is prognostic in the NDUFB4 diploid population. **(B)** Representative analysis of deconvoluted immune infiltrates within lung squamous cell carcinoma (LUSC) PanCancer TCGA dataset revealed negative correlation between NDUFB5 amplification status (Diploid number; moderate gain of copies; and amplification) and CD8$^{+}$ T cell abundance. **C)** Representative immunofluorescence analysis of CD8$^{+}$ T cell infiltration (shown in *green*) into CMT167 Parent or NDUFB5 OE tumors grown in C57BL/6 mice. Cropped from original images taken at 10x magnification using EVOS M7000 Imaging System. **(D)** Quantification of CD8$^{+}$ T cell abundance in CMT167 parent and NDUFB5 OE tumors (n=5); values shown as foci/field of view (FOV). Quantified from 20 FOVs/animal.
